# RBAi Scholar: Semantic and hybrid biomedical literature search for researchers and AI assistants

**DOI:** 10.64898/2026.11.26.769466

**Authors:** Hariom Chaudhary, Rimpy Kaur Chowhan

## Abstract

**Motivation:** A keyword search for *mitochondrial dysfunction* can miss a paper that says *impaired mitochondrial function*, so an empty result does not show that no such paper exists. Search by meaning addresses this mismatch, but it blurs exact strings such as gene symbols.

**Results:** We present RBAi Scholar, which lets researchers and AI assistants search a growing corpus of biomedical abstracts by meaning (semantic search), by keyword, or by meaning among the papers that contain a required term (hybrid search). We tested it on a fixed version of 11.1 million records. For 500 questions worded unlike their target papers, semantic search placed the target in the top ten for 65.0%, against 18.6% for LitSense 2.0 and 0.8% for PubMed. OpenAlex semantic search, documented as using GTE-Large-EN embeddings, reached 27.8%. It reached 45.8% after OpenAlex rebuilt it with Qwen3-Embedding-0.6B, the model RBAi Scholar uses. Semantic search also led on the 380 held-out targets, which played no part in its design. Given one required term from the paper, hybrid search reached 93.9% on 328 held-out targets, against 42.1% for LitSense 2.0 with the same term. For 86 queries made of a paper’s rarest identifier, such as a gene symbol, and two title words, keyword search reached 79.1%, against 34.9% for semantic search. When a blinded language-model judge graded the relevance of each system’s top ten results, semantic search returned the most relevant results (nDCG@10 0.651 against 0.592 for OpenAlex semantic search with Qwen3-Embedding-0.6B). Hybrid search ranked the target first more than twice as often as semantic search (50.0% against 23.3%). When we re-ran the searches for target papers on two cheaper index implementations and on the live service at 24.6 million records, every comparison kept its direction.

**Availability and implementation:** Web, REST API and Model Context Protocol server at https://thedreamlabinnovations.com/rbai-scholar.

Benchmark data and code: https://github.com/Insightwithhari/rbai-scholar.

**Graphical abstract:** 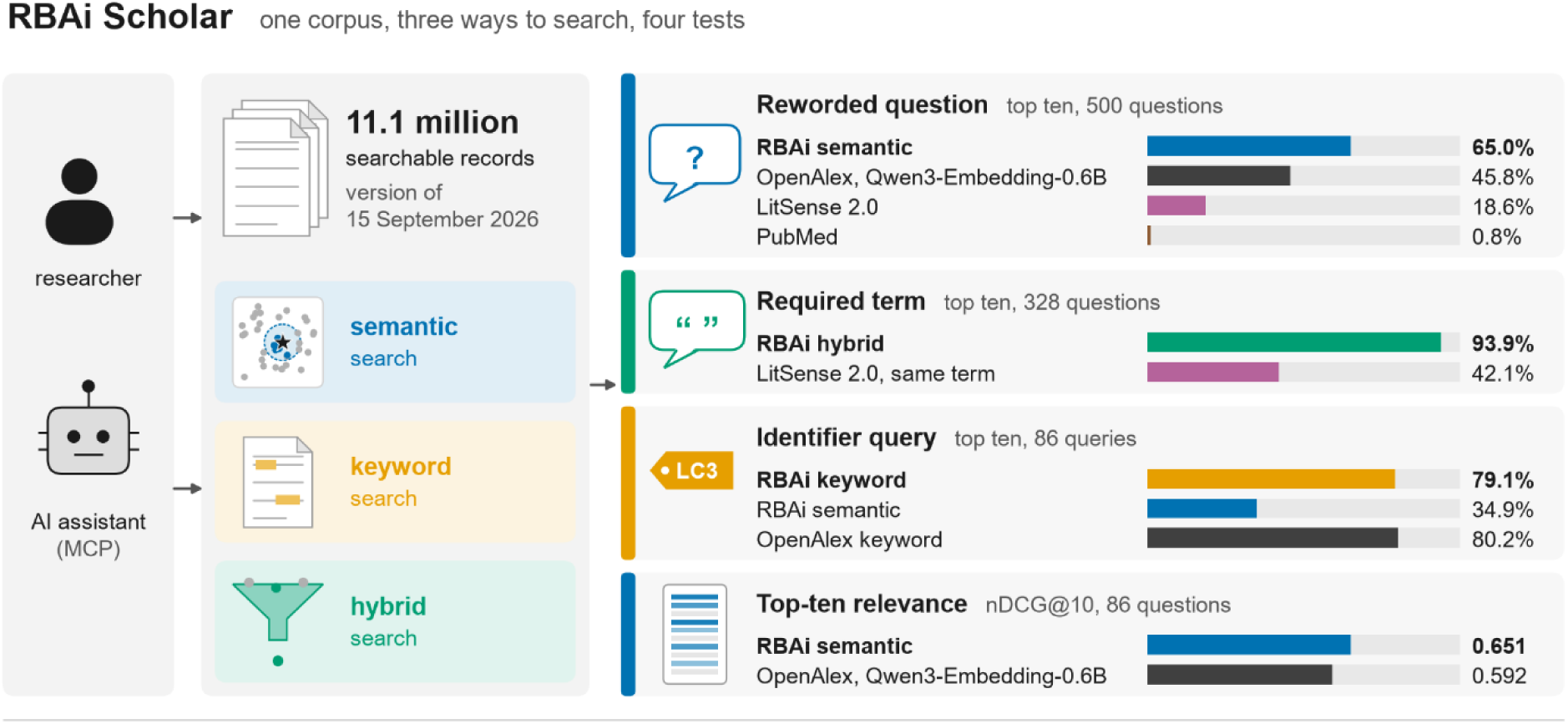

## 1 Introduction

Keyword search compares the words a searcher types with the words an author wrote. A researcher who types *mitochondrial dysfunction* can miss a paper that says *impaired mitochondrial function*, and nothing on the page shows the miss. PubMed adds the subject heading Mitochondrial Diseases to this search automatically. Even so, on 24 November 2026 it did not return 650 of the 1,582 PubMed records whose title or abstract says *impaired mitochondrial function*. An empty result looks the same whether no such paper exists or the paper uses other words. Trained searchers work around this with Medical Subject Headings and long Boolean queries. Many early-career researchers have not had that training, and the AI assistants that now search on their behalf send fluent sentences rather than carefully built query strings.

Semantic search addresses this mismatch. A language model maps each text to an *embedding*, a list of numbers placed so that texts with similar meaning lie close together. Papers are then ranked by closeness of meaning rather than by shared words. In biomedicine, MedCPT (Jin et al., 2023) trained a retriever and a re-ranker on 255 million query–article pairs from PubMed search logs. LitSense 2.0 (Yeganova et al., 2025) applies MedCPT to 38 million PubMed abstracts and 6.6 million PubMed Central full texts. It selects candidates by keyword matching and re-ranks the top 100 by meaning, so a paper that the keyword step misses cannot be recovered. Newer general-purpose embedding models, such as Qwen3-Embedding, score higher than earlier models on standard benchmarks (Zhang *et al*., 2025). Searchers, however, bring several needs: finding a paper from a description in other words, finding a paper they partly remember, finding it from an identifier, and surveying a topic.

These needs favour different kinds of matching. Meaning-based search can find a paper from a question that shares none of its words, but an embedding blurs exact strings such as a gene symbol or a trial number (Sciavolino et al., 2021; Xu et al., 2025), which keyword search matches exactly. A searcher who remembers one distinctive term needs both: the term to narrow the field, and meaning to order what remains. Our thesis is that no single mode serves every need, so a literature search service should offer each mode, and an evaluation should test each one on the need it serves.

We therefore built RBAi Scholar, a search service for biomedical abstracts with three modes. Semantic search ranks papers by meaning, keyword search by the words they contain, and hybrid search ranks by meaning only the papers that contain a required term. We tested it against PubMed, OpenAlex and LitSense 2.0 with an open benchmark of four tests, one for each of the needs above. Three tests ask whether a system finds one target paper from a question worded unlike the paper, from the same question with one term from the paper, or from a short query built around an identifier. The fourth test asks how relevant the top ten results are. It found the most target papers from reworded questions, and it returned the most relevant top ten. Hybrid search found the most target papers when the question carried a term from the paper. Keyword search found more than twice as many target papers from identifier queries as semantic search did. When we repeated the searches for target papers on cheaper indexes and on the live RBAi scholar service, which had grown to 24.6 million records, every comparison kept its direction.

## 2 Materials and methods

### 2.1 The corpus and the three search modes

We built the corpus from OpenAlex (Priem et al., 2022). It holds journal articles that have an abstract and whose OpenAlex primary topic lies in biomedicine or the adjacent life sciences. We removed duplicate records by DOI. We embedded each record’s title and abstract with Qwen3-Embedding-0.6B (Zhang et al., 2025) as unit-length vectors of 1,024 numbers. Then tested one fixed version of the corpus, the version of 15 September 2026. It holds 11,149,057 searchable records. All three retrieval modes search the same records (Fig. 1a). Semantic search puts the short task instruction that the model was trained with in front of the query, and embeds the result with the same model. It then re-ranks the 200 records nearest in meaning by this score:

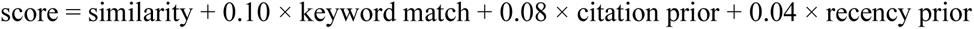

Similarity is the cosine similarity of the two embeddings. The citation prior grows with the logarithm of the citation count and stops growing at 339 citations, the 99th percentile of the corpus, so that a few highly cited papers cannot dominate. The recency prior rises evenly over 60 years (Supplementary Section S1). Keyword search uses a full-text index with familiar syntax (all words required by default, AND, OR, NOT and quoted phrases) and ranks by BM25 (Robertson and Zaragoza, 2009) plus the same two priors. In hybrid search, the searcher writes a question and puts one term in quotation marks, for example *what stabilises “R175H” mutant p53*. Only papers that contain the term are eligible, and the question orders them as in semantic search. Hybrid search declines a term present in more than 20,000 records, because such a term would not narrow the search.

**Fig. 1.**
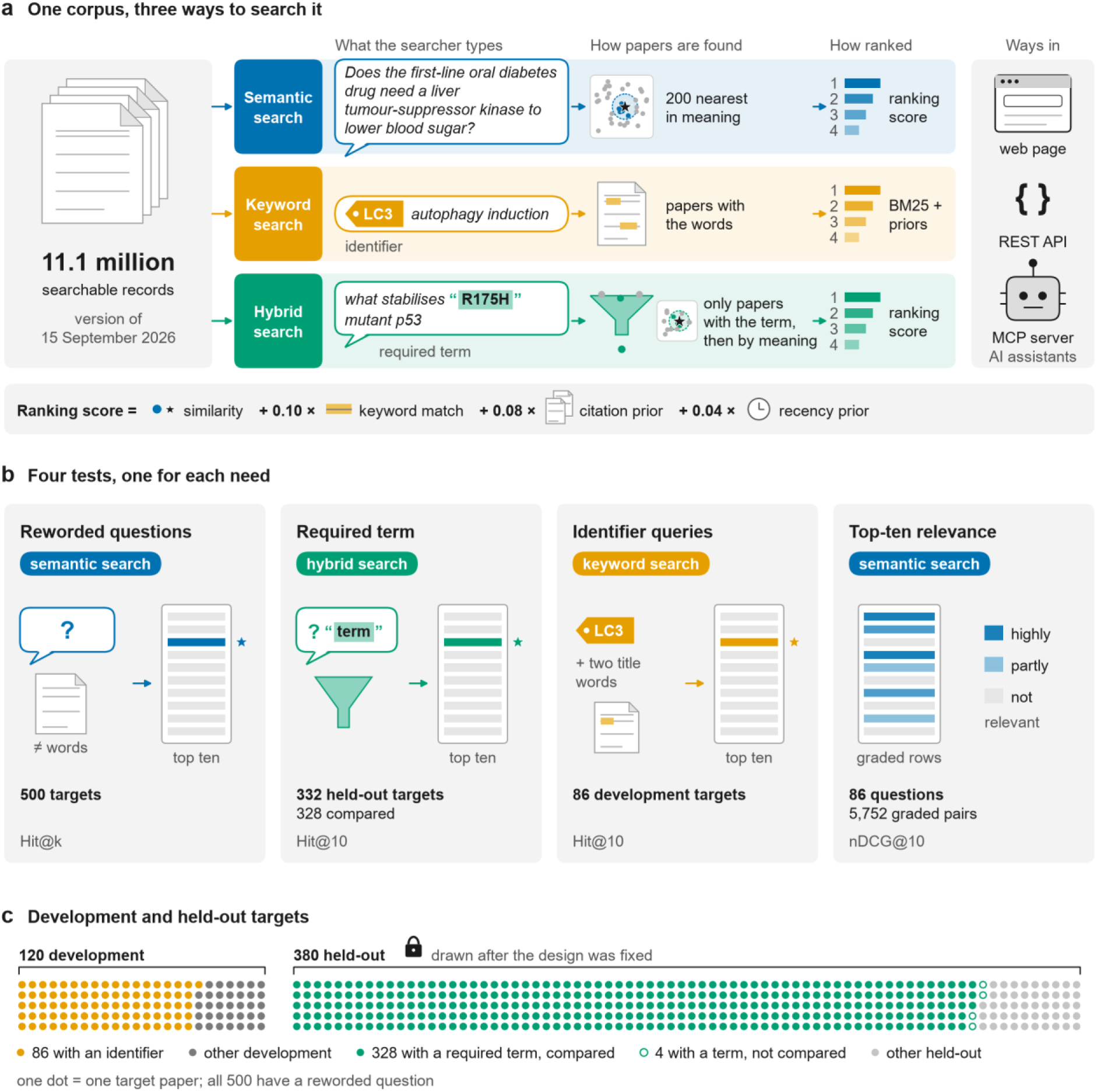
RBAi Scholar searches one corpus in three ways, and four tests measure each way on the need it serves.

RBAi Scholar is available through a web page, a REST API and a Model Context Protocol (MCP) server (Model Context Protocol, 2025), which lets AI assistants call it directly. The MCP server offers semantic and keyword search as separate tools, and a quoted term in a semantic search gives hybrid search.

### 2.2 The benchmark

The benchmark has four tests (Fig. 1b), and it keeps the targets used to develop the service apart from those used to evaluate it (Fig. 1c). Every question, query and per-question result is in Supplementary Data 1.

**Targets.** We drew 500 target papers from the corpus, 100 from each of five citation bands (0–9, 10–49, 50–199, 200–999 and ≥1,000 citations), so that rarely cited papers, about half of the corpus, are properly represented. The first 120 (24 per band) are development targets, used for every design experiment (Supplementary Table S1). The other 380 (76 per band) are held-out targets, drawn after the configuration had been fixed and used only to measure it.

**Reworded questions.** For each target, a language model (Claude Opus 5, Anthropic) wrote one question that the paper answers. It worked from the title and abstract, and we told it to avoid the paper’s distinctive wording. On average, 39.2% of the words in a development question and 40.4% of those in a held-out question appear anywhere in the target abstract, and no question shares a phrase longer than five words with it (Supplementary Data 1). Our tests do not use text copied from the abstracts, because copied text rewards word matching. In a check on the 120 development targets, queries made of each abstract’s first 350 characters at least doubled the top-ten score of each of the three systems tested (Supplementary Table S1).

**Required terms.** Hybrid search needs a quoted term. For each held-out target, the language model that wrote the questions chose one term that a searcher would plausibly type with the question. The term had to appear word for word in the title or abstract, and the model placed it where it reads naturally in the question. We call this the relevance rule. It found a term for 332 of the 380 targets. Hybrid search answered 328 of these 332 searches, and comparisons between systems use those 328 (Supplementary Table S3). Counting the other 4 as misses would lower hybrid search’s top-ten score from 93.9% to 92.8%.

**Identifier queries.** For the 86 development targets that contain an identifier, such as a gene symbol, a variant or an accession, we built the short kind of query a searcher types. A fixed rule took the target’s rarest identifier and added two title words, for example *LC3 autophagy induction*. Earlier studies built simulated searches in the same way (Azzopardi et al., 2007). Every keyword system received the same strings.

**Relevance of the top ten.** For the same 86 targets, every system searched with the input its mode is built for. Semantic searches received the reworded question. Keyword searches received the identifier query, and hybrid search received the question with the rarest identifier in quotation marks. OpenAlex, which has no hybrid mode, also received the question with the identifier appended, which is the text hybrid search received without the quotation marks. We pooled every paper in every top ten, including the papers from OpenAlex’s Qwen3-Embedding-0.6B run (Section 2.3). The pool held 5,752 question– paper pairs. A language model (Claude Haiku 5.5, Anthropic) graded every pair. It saw only the question, the title and the abstract, so it never knew which system had returned a paper or at what rank. It wrote a reason for every grade under a written rubric (Supplementary Section S2). Grade 0 means a different subject, 1 the same general area only, 2 a paper that answers part of the question, and 3 one that answers exactly what was asked. Grades 2 and 3 count as relevant. Earlier studies found that language-model relevance labels agree closely with human assessors and order systems as human labels do (Thomas et al., 2024; Upadhyay et al., 2024).

### 2.3 Comparators

We sent each reworded question unchanged to OpenAlex semantic search (the search.semantic parameter of its /works endpoint), to PubMed through the E-utilities with its relevance ranking, Best Match (Fiorini et al., 2018), and to LitSense 2.0 passage search with MedCPT re-ranking switched on (Yeganova et al., 2025). LitSense 2.0 returns passages, so we mapped each passage to its paper by PubMed identifier, and each paper kept the rank of its best passage. We queried the comparators in September 2026, when OpenAlex’s documentation said that its semantic search used GTE-Large-EN embeddings. On 23 September 2026, OpenAlex announced that it had rebuilt its semantic search with Qwen3-Embedding-0.6B, the model RBAi Scholar uses (Priem, 2026). We therefore sent OpenAlex the 500 questions and the 86 searches of the relevance test again in November 2026, with the same code and settings. We name the two OpenAlex runs after these two models, as the GTE-Large-EN run and the Qwen3-Embedding-0.6B run. OpenAlex’s keyword search uses no embeddings, and the results below use its lists from the Qwen3-Embedding-0.6B run. OpenAlex holds all 500 targets and PubMed holds 499 of them, so a miss by either system almost always comes from its ranking rather than from a missing paper.

### 2.4 Measures and statistics

*Hit@k* is the share of questions whose target appears in the first *k* results. For the relevance of the top ten we report two measures. *P@shown* is the share of the returned results that were graded relevant. *nDCG@10* (Järvelin and Kekäläinen, 2002) gives more credit to more relevant papers and to higher places, and it runs from 0 to 1. Confidence intervals (CI) are 95% bootstrap intervals over questions (10,000 resamples). Paired comparisons of Hit@k use the exact McNemar test, which counts the questions on which only one of the two systems found the target. Differences in nDCG@10 and P@shown use a paired permutation test with 20,000 sign flips, so the smallest value it can report is *P* < 5 × 10⁻⁵. Where several tests answer one question, *P* values are corrected for the number of tests by Holm’s method and marked Holm-adjusted. A declined or unanswered search counts as a miss, except in the required-term comparison, which uses the 328 questions that every system answered.

### 2.5 Index implementations and the live service

Semantic search must find the nearest of 11.1 million embeddings for every query. Approximate nearest-neighbour indexes make this fast by giving up a little recall, and they differ greatly in the memory they need. We measured every result on a reference implementation, an HNSW graph (Malkov and Yashunin, 2020) in PostgreSQL held in 87.4 GB of memory. We then re-ran every published search on two cheaper implementations. One is a disk-resident DiskANN graph (Subramanya et al., 2019) of 5.0 GB. The other is a serverless deployment, which runs on Cloudflare’s managed services without a server of our own, with Vectorize for the embeddings and D1 databases for the records. The live service is the serverless deployment, grown to 24.6 million records.

## 3 Results

### 3.1 Meaning-based search finds papers from reworded questions that word-matching systems miss

The first test asked whether a search engine can find a particular paper when the question describes its findings in other words. On the 500 reworded questions, semantic search ranked the target first for 32.8% of the questions and placed it in the top ten for 65.0% (Fig. 2a; Supplementary Table S2). OpenAlex semantic search with GTE-Large-EN ranked the target first for 9.0% and placed it in the top ten for 27.8%. The same figures were 6.0% and 18.6% for LitSense 2.0, and 0.6% and 0.8% for PubMed. When we compared RBAi Scholar with LitSense 2.0 question by question, RBAi Scholar placed 241 targets in its top ten that LitSense 2.0 missed, and LitSense 2.0 placed 9 that RBAi Scholar missed (exact McNemar test, *P* < 10⁻⁶). RBAi Scholar’s top-ten score was the highest in every citation band, including the papers cited 0 to 9 times, which make up about half of the corpus (50.6%). For these papers, RBAi Scholar placed 54% of the targets in the top ten, against 12% for OpenAlex with GTE-Large-EN and 21% for LitSense 2.0 (Fig. 2b).

**Fig. 2.**
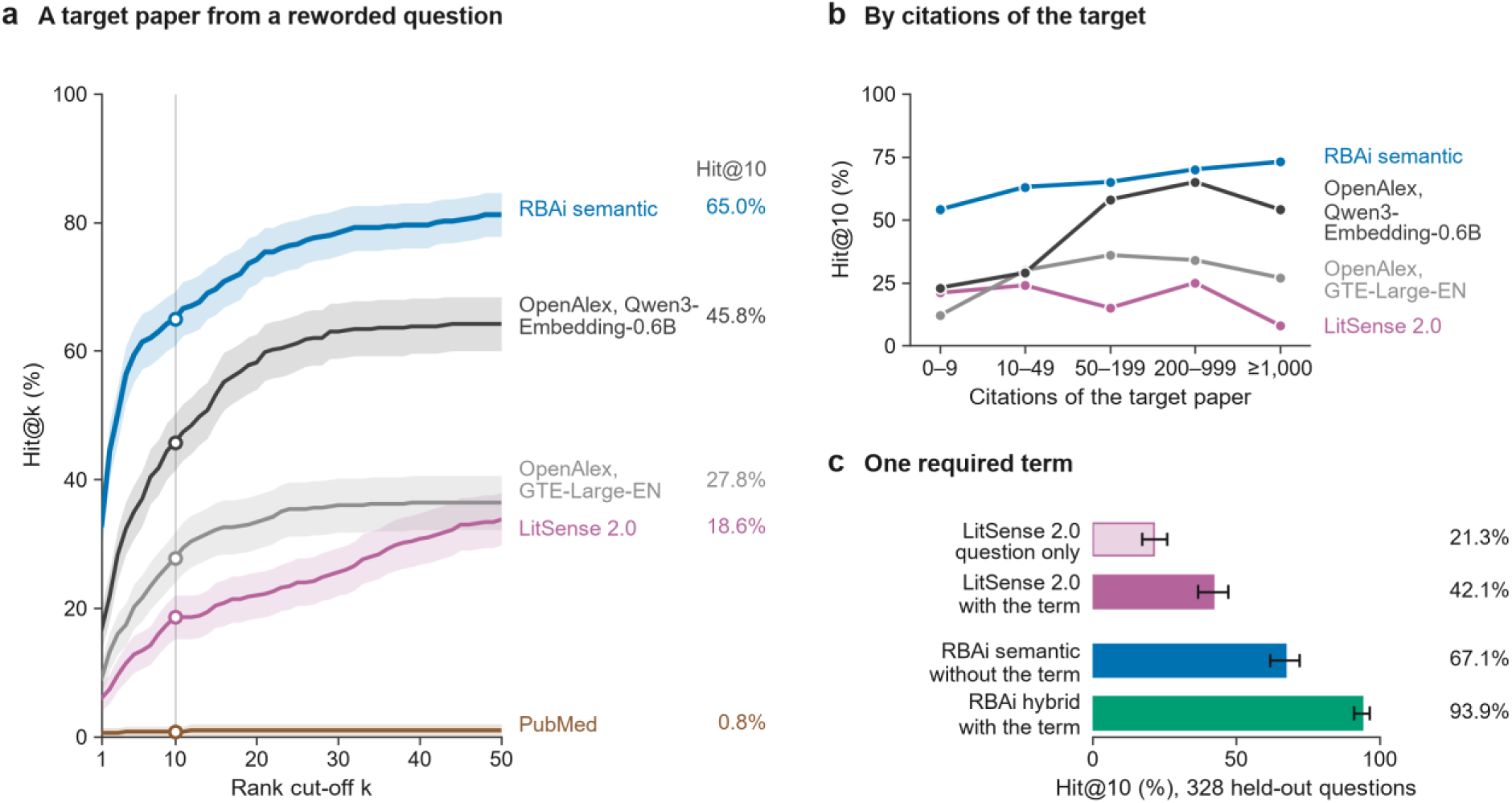
RBAi Scholar finds more target papers than OpenAlex, LitSense 2.0 and PubMed. (a) The share of the 500 reworded questions whose target paper is in the top *k* results (Hit@k), for *k* from 1 to 50, with *k* = 10 marked by the vertical line. (b) Hit@10 in each citation band, 100 targets per band. PubMed, which found at most 2 targets in any band, is not shown. (c) Hit@10 on the 328 held-out questions with a required term, for LitSense 2.0 without and with the term, RBAi Scholar’s semantic search without the term and its hybrid search with it. Shaded bands and error bars are 95% bootstrap intervals. Data: Supplementary Data 1.

The way the other systems find candidates helps explain the gap. LitSense 2.0 first selects candidates by matching words and only then re-ranks the top 100 by meaning, so a paper that the word-matching step misses cannot come back. PubMed combines the words of a query with AND, so it returns only records that match all of them. For 466 of the 500 questions, it returned no papers at all. We also shortened each of the 120 development questions to its four longest content words, joined by AND. The shortened queries cut the questions with no result from 104 to 34, but they raised PubMed’s top-ten score only from 1.7% to 2.5% (Supplementary Table S1). These figures show the limits of word matching on questions written in other words. They do not measure how well PubMed serves the structured Boolean searches it was built for.

Could our design choices have been fitted to the development targets? On the 380 held-out targets, drawn only after every design choice had been fixed, semantic search placed 67.6% in the top ten, against 56.7% on the 120 development targets (Supplementary Table S2). LitSense 2.0 also scored higher on them (20.8% against 11.7%). The four systems ranked in the same order in both sets.

Was RBAi Scholar ahead only because of its embedding model? OpenAlex has since rebuilt its semantic search with Qwen3-Embedding-0.6B, the model RBAi Scholar uses (Section 2.3). When we sent OpenAlex the same 500 questions after this change, it placed the target in the top ten for 45.8%, up from 27.8%, and for 48.2% of the held-out targets. RBAi Scholar remained about 19 percentage points ahead, at 65.0% overall and 67.6% on the held-out targets. When we compared the two question by question, 102 targets were in RBAi Scholar’s top ten but not in OpenAlex’s, and 6 were the other way round (*P* < 10⁻⁶). OpenAlex gained most on highly cited papers. Its top-ten score rose from 34% to 65% for papers with 200 to 999 citations, but only from 12% to 23% for those with 0 to 9 (Fig. 2b). For targets with at least 1,000 citations, OpenAlex now ranked the target first more often than RBAi Scholar did (43% against 34%), although this difference was not statistically significant (14 targets against 5, *P* = 0.064). RBAi Scholar still placed more of these targets in the top ten (73% against 54%).

Both systems now use the same embedding model, so the remaining gap must come from other differences between them. OpenAlex embeds each work’s title, abstract and venue name, and among close matches it gives works with more citations a small boost (OpenAlex, 2026). That boost may help explain its strength on highly cited papers. RBAi Scholar adds to each question the task instruction the model was trained with. On the development questions, the instruction alone raised the top-ten score from 44.2% to 58.3% (Supplementary Table S1), and OpenAlex’s documentation does not mention one.

OpenAlex also searches hundreds of millions of works from every field of research and of every type, so papers from other fields, and records without an abstract, may compete for its top places. We did not test which of these differences matters most.

In short, both meaning-based engines found more target papers than the two systems that start by matching words, and RBAi Scholar found the most, even after OpenAlex adopted the same embedding model.

### 3.2 A required term helps, and ranking by meaning within it helps most

A searcher who has read a paper before may remember one distinctive term from it. Does requiring that term help, and which system uses it best? We tested this on 328 held-out targets with a term chosen by the relevance rule (Section 2.2). Hybrid search receives the term, so we ran LitSense 2.0 twice, once with the question alone and once with exactly the text that hybrid search received, including the term. The term almost doubled LitSense 2.0’s top-ten score, from 21.3% to 42.1% (Fig. 2c). Given the same text, hybrid search placed 93.9% of the targets in the top ten and ranked 71.0% first, against 17.4% for LitSense 2.0. When we compared the two question by question, 172 targets were in hybrid search’s top ten but not in LitSense 2.0’s, and 2 were the other way round (*P* < 10⁻⁶; Supplementary Table S3). Even semantic search, which never saw the term, placed 67.1% of the targets in the top ten, more than LitSense 2.0 did with it. Because each term was taken from its target paper, these figures describe a searcher who remembers a term that the authors actually used.

A required term narrows the search for any system, and it helped most when every paper containing the term was ranked by meaning, as hybrid search does.

### 3.3 Identifiers need keyword search, which finds papers that semantic search misses

Not every search is a question. Researchers also search with short strings, such as a gene symbol or an accession number with a word or two of context. On the 86 identifier queries (Section 2.2), keyword search placed the target in the top ten for 79.1% (68 of 86), and semantic search did so for 34.9% (30 of 86; Fig. 3a, Supplementary Table S3). This fits earlier reports that embeddings blur exact strings such as gene symbols (Sciavolino et al., 2021; Xu et al., 2025). Quoting the query’s rarest word, which was the identifier itself in 78 of the 86 queries, recovered most of this difference. Hybrid search then reached 70.9% (61 of 86). Compared with semantic search without quotation marks, it gained 32 targets and lost 1 (*P* < 10⁻⁶).

**Fig. 3.**
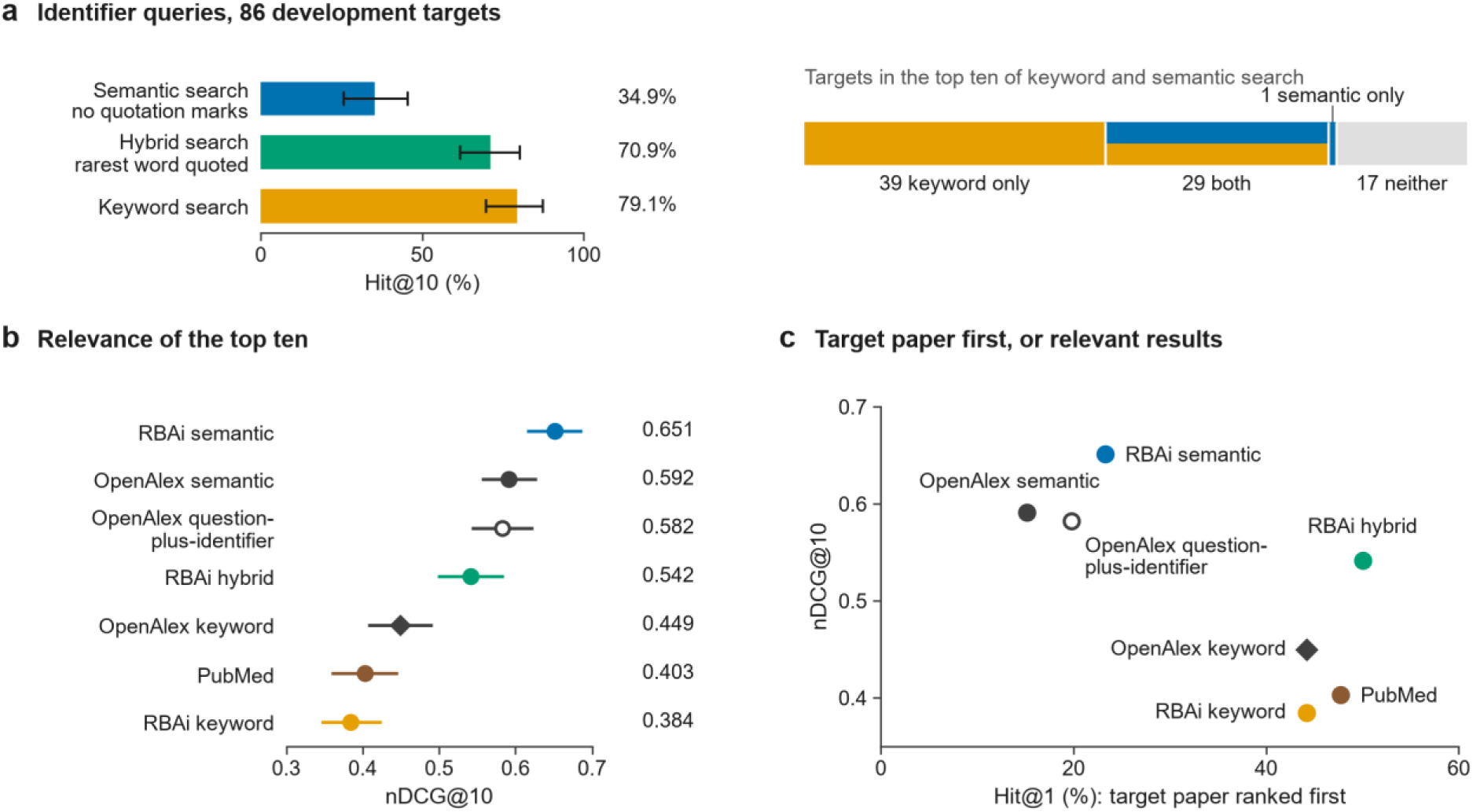
Keyword search finds target papers from identifier queries, and semantic search returns the most relevant top ten. (a) Hit@10 on the 86 identifier queries for semantic search without quotation marks, hybrid search with the rarest word quoted, and keyword search. The strip shows which targets keyword search and semantic search placed in their top ten. (b) nDCG@10 of the top ten on the 86 graded questions, for RBAi Scholar’s three modes, OpenAlex’s Qwen3-Embedding-0.6B run and PubMed. (c) The share of questions with the target paper ranked first (Hit@1) against nDCG@10, for the same systems. Error bars are 95% bootstrap intervals. Data: Supplementary Data 1.

Keyword search also reached papers that semantic search could not. It found 29 of the 30 targets that semantic search found, and 39 more. Neither mode found the other 17. In a development experiment, we also merged the two ranked lists into one by reciprocal rank fusion (Cormack et al., 2009). The merged list kept keyword search’s top-ten score, but it ranked the target first only a third as often (14.0% against 43.0%; Supplementary Table S1). Semantic search ranks these queries poorly, so merging pushed the correct answers down. RBAi Scholar therefore keeps the modes separate and lets the searcher, or the AI assistant, choose the one that suits the query.

### 3.4 Semantic search returns the most relevant top ten, and hybrid search ranks the target paper first

The tests so far score one paper and ignore the other nine results. Are those results worth reading? For the 86 targets of Section 3.3, a blinded judge graded every paper in every system’s top ten, with OpenAlex represented by its Qwen3-Embedding-0.6B run (Section 2.2). We summarise each top ten by nDCG@10 (Section 2.4) and by the number of relevant papers in it.

RBAi Scholar’s semantic search returned the most relevant top ten of any system tested. Its nDCG@10 was 0.651 (95% CI 0.615–0.687), and on average 7.43 of its ten results were relevant (Fig. 3b; Supplementary Table S4). OpenAlex semantic search scored 0.592, and its question-plus-identifier variant scored 0.582. Both scored lower than RBAi Scholar’s semantic search (Holm-adjusted *P* < 0.001 and *P* = 0.004).

Hybrid search showed a trade-off between finding the target paper and returning other relevant papers (Fig. 3c). It ranked the target paper first for half of the questions (50.0%), against 23.3% for semantic search, which received the same question without the quoted term. Its top ten, however, was less relevant than semantic search’s (nDCG@10 0.542 against 0.651; Holm-adjusted *P* = 0.002). The required term lifts the target paper to the top, but it probably also lets in papers that share the term without sharing the topic.

Our keyword search is selective. It returned fewer results than OpenAlex keyword search (5.83 against 9.43 on average), and more of them were relevant (P@shown 0.642 against 0.411; Holm-adjusted *P* < 0.001). Over the whole top ten, however, it trailed OpenAlex’s keyword search (nDCG@10 0.384 against 0.449; Holm-adjusted *P* = 0.0006). Part of this gap probably comes from the measure itself, which counts every empty place in the top ten against the system that returns fewer results. Both found the target paper in the top ten about equally often (79.1% for ours and 80.2% for OpenAlex). Only 5 targets were found by ours alone and 6 by OpenAlex alone (*P* = 1).

### 3.5 Searches for target papers gave similar results on cheaper indexes and on a corpus 2.2 times larger

We re-ran every published search, unchanged, on the DiskANN index and on the serverless deployment (Section 2.5; Supplementary Table S5). The three implementations gave nearly the same results. On the 500 reworded questions, semantic search placed the target in the top ten for 65.0% on the reference, 65.2% on DiskANN and 66.0% on the serverless deployment. Keyword search placed 79.1%, 79.1% and 82.6% of the identifier targets in the top ten on the same three implementations. The serverless deployment has its own full-text index, which found 3 more targets than the reference did and lost none. Only one set of searches changed with the nearest-neighbour index. When we sent the identifier queries to semantic search without quotation marks, it placed 34.9% of the targets in the top ten on the reference, 39.5% on DiskANN and 26.7% on the serverless deployment. With the query’s rarest word in quotation marks, hybrid search placed 70.9% in the top ten on all three.

By 16 November 2026 the live service held 24.6 million records, 2.2 times as many as the tested corpus. It used the same model, ranking and weights. When we sent it the same searches for target papers, every comparison kept its direction. Semantic search placed 63.8% of the 500 targets in the top ten, against 65.0% on the tested corpus. Hybrid search placed 93.3% of the 328 required-term targets in the top ten, against 93.9%. The reference and the serverless deployment differ slightly on the tested corpus, so we measured the cost of the larger corpus on the serverless deployment alone. Its top-ten score fell from 66.0% to 63.8%. Two targets entered the top ten and 13 left it (*P* = 0.007). Its share of targets ranked first also fell, from 33.0% to 30.4%. One target moved up to first place and 14 dropped from it (*P* = 0.001), probably because more papers now compete for each top place.

## 4 Discussion

The three modes do not form a ranking from best to worst. Each answers a different need (Fig. 3c):

- To survey a topic, use semantic search. It returned the most relevant top ten of any system tested.
- To find a paper you partly remember, use hybrid search and quote a term you remember from the paper. In the held-out test it placed 93.9% of the targets in the top ten, and on the graded questions it ranked the target paper first half of the time.
- For identifiers, exact phrases and Boolean logic, use keyword search. It found the target of an identifier query more than twice as often as semantic search.

An AI assistant faces the same choice on every request. Through the MCP server it sees semantic and keyword search as separate tools, each described by the kind of query it serves, so it can send fluent questions to semantic search, identifiers to keyword search, and a quoted term when the target is one specific paper. Because the results carry abstracts and PubMed identifiers, the assistant can read and cite what it finds.

## 5 Conclusion

RBAi Scholar lets researchers and AI assistants search biomedical abstracts in the way each need requires: by meaning to find a paper from a question in other words or to survey a topic, by keyword for identifiers, and by meaning within a quoted term to find a paper they partly remember. On reworded questions it found target papers more often than OpenAlex, LitSense 2.0 and PubMed, including on held-out targets and after OpenAlex adopted the same embedding model. Its lead in finding target papers held on three index implementations and on a corpus 2.2 times larger. The next step is re-ranking. In our tests on the reworded questions, MedCPT’s cross-encoder (Jin *et al*., 2023) raised the share of targets ranked first from 32.8% to 48.6% (Supplementary Table S6). The open benchmark is released so that others can test their systems on the needs their users have.

## Author contributions

H.C. built the resource, designed and ran the evaluation, analysed the results and wrote the manuscript.

R.K.C. guided the biomedical framing and revised the manuscript. Both authors approved the final version

## Use of AI tools

Large language models (Anthropic Claude Opus 5.5) helped draft and edit this text and write analysis code. The authors checked every reported number against the released data files, All contents were verified. Claude Opus 5 (Anthropic) wrote every benchmark question from the target’s title and abstract. We told it to avoid the paper’s distinctive wording, and we measured the overlap with the abstract afterwards. The same model chose each relevance-rule term under written criteria and placed it in the question. Claude Haiku 5.5 (Anthropic), acting as a blinded judge, produced the relevance grades under the written rubric in Supplementary Section S2 (5,752 grades and a re-grading of 217). Every grade and its reason is released.

## Data availability

Supplementary Data 1 lists every target, question and query with each system’s result for each question. It also lists each result file with its date. The full benchmark is released at https://github.com/Insightwithhari/rbai-scholar. It contains every query as sent, per-question ranks for every system, every grade with its reason, the lists from OpenAlex’s Qwen3-Embedding-0.6B run, and the re-runs on the other two index implementations and on the live service. Running node benchmark/score.mjs (Node.js 18 or later, no installation) recomputes the published results from these files without contacting the live service. The search service is accessible at https://thedreamlabinnovations.com/rbai-scholar, through the REST endpoint https://scholar.thedreamlabinnovations.com/v1/corpus/search and the MCP endpoint https://scholar.thedreamlabinnovations.com/v1/mcp benchmark repository is available at https://github.com/Insightwithhari/rbai-scholar.

## Supplementary Information

Section S1 gives the ranking details that the main text leaves out. Section S2 gives the instruction that the relevance judge received. Tables S1 to S6 hold the design experiments and the full results behind the main text. Supplementary Data 1 is an Excel workbook that lists every target, question and query, with each system’s result for each question. Files named in code font are in the benchmark folder of https://github.com/Insightwithhari/rbai-scholar.

### S1 Ranking details

**Candidates and embeddings.** Semantic search and hybrid search each re-rank a pool of 200 candidates, whatever number of results is requested. We joined each record’s title and abstract as “title. abstract” and embedded the text in half precision. Only queries receive the task instruction, because the model was trained that way (Zhang *et al*., 2025).

**The three components beside similarity.** The keyword match scores each candidate against a text query that requires every word of the question. The search then scales these scores within the pool of 200. The citation prior is min(1, ln(1 + citations) / ln(1 + 339)). The recency prior is 0.5 when the year is missing.

**Keyword ranking.** Keyword search ranks by BM25 (Robertson and Zaragoza, 2009) with *k*₁ = 1.2 and *b* = 0.75. It counts the word statistics over all searchable records. It scales the BM25 score to the range 0 to 1 and then adds the two priors. When more than 2,000 records match a query, keyword search scores only the 2,000 most cited of them. Every identifier query matched between 1 and 180 records, so keyword search scored every matching record.

**Quoted terms.** The search reduces the words of a quoted term to their stems, so that *mutants* also matches *mutant*. The words must appear together as a phrase. The search removes the quotation marks before it embeds the question, but the words stay. The term therefore adds to the meaning of the question as well as to the filter.

### S2 Relevance rubric

The judge, Claude Haiku 5.5 (Anthropic), received the instruction below with each batch. We quote its grade definitions and rules word for word. The rest of the instruction sets the output format. The released file JUDGING-PROTOCOL.md gives the same four grades.

> For every document, decide how well it answers its question, and give it one grade with a written reason. > > 3 Directly answers the question. The paper’s own subject and findings address exactly what was asked. A researcher would cite this as the answer. > > 2 Substantially relevant. Addresses the question and supplies a real part of the answer, or answers it for a closely related system, organism or condition. > > 1 Marginally relevant. Same general subject area. Useful background or context, but does not address what was asked. > > 0 Not relevant. A different subject, or shares only generic vocabulary with the question. > > Rules > - Judge each document only from its question, its title and its abstract. Where abstract_available is false, judge from the title alone. > - Keep the question in mind and grade its documents against the same standard. The documents are listed in random order; the order says nothing about them. > - Every grade needs a reason of at least 25 characters naming what in the title or abstract justified it. > - Grade every document of the batch exactly once: no document skipped, none added. > - Use only the batch file and these instructions. Do not open other files, look anything up, or use any other source.

nDCG@10 gives a paper of grade *g* a gain of 2*^g^* − 1 and divides that gain by log_2_(rank + 1), so that lower places count less (Järvelin and Kekäläinen, 2002). It adds these values up over the top ten and divides the sum by the largest sum that the question’s graded papers allow. A paper listed twice in one top ten counts once.

### Supplementary tables

**Table S1.** Design experiments on the 120 development targets. Most of these experiments ran on development versions of the service, so their values differ from those of the tested configuration. Hit@10 is the share of targets placed in the top ten, and Hit@1 the share ranked first. The column *Change in the top ten* compares a setting with the setting it names, target by target. For example, compared with the plain query, the task instruction brought 19 targets into the top ten (gained) and dropped 2 (lost), and the exact McNemar test gives the *P* value. Data: data/ablation-prefix-depth.jsonl, data/known-item-240.jsonl, data/identifier-86.jsonl, data/fusion-identifier-86.jsonl, data/marked-term-quality.jsonl, data/hybrid-86-paired.jsonl, data/placement-86.jsonl.

| Experiment and setting | Hit@10 | Hit@1 | Change in the top ten | Outcome |
|---|---|---|---|---|
| **Semantic search, 120 reworded questions** |  |  |  |  |
| Plain query, pool of 40 candidates | 44.2% | 15.8% |  |  |
| Pool of 200 candidates | 47.5% | 14.2% | 8 gained and 4 lost against the plain query, *P* = 0.39 |  |
| Task instruction | 58.3% | 21.7% | 19 gained and 2 lost against the plain query, Holm-adjusted *P* = 0.00066 |  |
| Both | 60.0% | 23.3% | 21 gained and 2 lost against the plain query, *P* = 0.0001 | adopted |
| **Copied passages and reworded questions, development version of the service** |  |  |  |  |
| RBAi Scholar, copied passages | 89.2% |  |  |  |
| RBAi Scholar, reworded questions | 44.2% |  |  |  |
| OpenAlex semantic search with GTE-Large-EN, copied passages | 54.2% |  |  |  |
| OpenAlex semantic search with GTE-Large-EN, reworded questions | 26.7% |  |  |  |
| PubMed, copied passages | 23.3% |  |  |  |
| PubMed, reworded questions | 1.7% |  |  |  |
| **PubMed with shortened queries** |  |  |  |  |
| Questions as typed, 104 of them with no result | 1.7% |  |  |  |
| Four longest content words joined by AND, 34 with no result | 2.5% |  |  |  |
| **86 identifier queries** |  |  |  |  |
| Semantic search, similarity alone | 17.4% | 7.0% |  |  |
| Keyword search, built-in ranking | 70.9% | 41.9% | 49 gained and 3 lost against semantic search, *P* < 10<sup>−6</sup> |  |
| Keyword search, BM25 | 75.6% | 43.0% | 6 gained and 2 lost against built-in ranking, *P* = 0.29 | adopted |
| BM25 merged with semantic results by reciprocal rank fusion (Cormack *et al.*, 2009) | 75.6% | 14.0% |  | not adopted |
| **Quoting the rarest word of 118 reworded questions** |  |  |  |  |
| Not quoted | 56.8% | 25.4% |  |  |
| Quoted | 16.1% | 10.2% | 1 gained and 49 lost against not quoted, *P* < 10<sup>−6</sup> | not adopted |
| **Which term hybrid search quotes, 86 development targets** |  |  |  |  |
| Relevance rule | 87.2% | 60.5% |  | adopted |
| Rarest-identifier rule | 76.7% | 50.0% | 3 gained and 12 lost against the relevance rule, *P* = 0.035 |  |
| **Position of the quoted term, 86 graded questions** |  |  |  |  |
| Five positions in the question | 74.4–76.7% |  |  |  |

**Table S2.** Finding target papers from 500 reworded questions. Hit@*k* is the share of questions whose target paper was among the first *k* results. We asked each system for 50 results, so every cutoff comes from one list. The last two columns compare RBAi Scholar with one other system, target by target. Each cell counts the targets that only one of the two systems placed within the cutoff, first for RBAi Scholar and then for the other system. For example, 241 targets were in RBAi Scholar’s top ten but not in LitSense 2.0’s, and 9 were in LitSense 2.0’s top ten but not in RBAi Scholar’s. Every one of these differences was statistically significant (exact McNemar test, *P* < 10⁻⁶). Data: Supplementary Data 1, sheet Reworded questions (500).

| Measure | RBAi Scholar semantic search | OpenAlex semantic search with GTE-Large-EN | OpenAlex semantic search with Qwen3-Embedding-0.6B | LitSense 2.0 | PubMed | Found by only one system: RBAi Scholar / LitSense 2.0 | Found by only one system: RBAi Scholar / OpenAlex with Qwen3-Embedding-0.6B |
|---|---|---|---|---|---|---|---|
| **All 500 questions** |  |  |  |  |  |  |  |
| Hit@1 | 32.8% | 9.0% | 16.6% | 6.0% | 0.6% | 142 / 8 | 101 / 20 |
| Hit@5 | 59.4% | 20.2% | 35.0% | 12.8% | 0.8% | 237 / 4 | 133 / 11 |
| Hit@10 | 65.0% | 27.8% | 45.8% | 18.6% | 0.8% | 241 / 9 | 102 / 6 |
| Hit@20 | 74.2% | 33.4% | 58.2% | 22.0% | 1.0% | 267 / 6 | 87 / 7 |
| Hit@50 | 81.2% | 36.4% | 64.2% | 33.8% | 1.0% | 245 / 8 | 92 / 7 |
| **Hit@10 by target set and citation band** |  |  |  |  |  |  |  |
| Development targets (120) | 56.7% | 26.7% | 38.3% | 11.7% | 1.7% |  |  |
| Held-out targets (380) | 67.6% | 28.2% | 48.2% | 20.8% | 0.5% |  |  |
| 0–9 citations (100) | 54.0% | 12.0% | 23.0% | 21.0% | 0.0% |  |  |
| 10–49 citations (100) | 63.0% | 30.0% | 29.0% | 24.0% | 1.0% |  |  |
| 50–199 citations (100) | 65.0% | 36.0% | 58.0% | 15.0% | 1.0% |  |  |
| 200–999 citations (100) | 70.0% | 34.0% | 65.0% | 25.0% | 0.0% |  |  |
| ≥1,000 citations (100) | 73.0% | 27.0% | 54.0% | 8.0% | 2.0% |  |  |

**Table S3.** Required terms and identifier queries. Hit@*k* is the share of questions whose target paper was among the first *k* results. (a) The 328 held-out questions that every system answered. The *P* values come from the exact McNemar test, adjusted by Holm’s method across the five cutoffs. (b) The 86 identifier queries. Hybrid search quotes the query’s rarest word. It declined the 2 terms present in more than 20,000 records, and these count as misses. Data: Supplementary Data 1, sheets Required term (332) and Identifier queries (86).

| System | Hit@1 | Hit@5 | Hit@10 | Hit@20 | Hit@50 |
|---|---|---|---|---|---|
| **(a) A required term, 328 held-out questions** |  |  |  |  |  |
| LitSense 2.0, question only | 7.6% | 14.3% | 21.3% | 25.3% | 39.6% |
| LitSense 2.0, question with the term | 17.4% | 34.5% | 42.1% | 51.2% | 67.7% |
| RBAi Scholar semantic search, no term | 33.8% | 61.0% | 67.1% | 75.6% | 82.6% |
| RBAi Scholar hybrid search* | 71.0% | 90.5% | 93.9% | 96.0% | 97.9% |
| Difference, hybrid search minus LitSense 2.0 with the term (percentage points) | +53.7 | +56.1 | +51.8 | +44.8 | +30.2 |
| *P*, hybrid search against LitSense 2.0 with the term | < 10<sup>−6</sup> | < 10<sup>−6</sup> | < 10<sup>−6</sup> | < 10<sup>−6</sup> | < 10<sup>−6</sup> |
| **(b) 86 identifier queries** |  |  |  |  |  |
| Semantic search, no quotation marks | 24.4% |  | 34.9% |  |  |
| Hybrid search, rarest word quoted | 39.5% |  | 70.9% |  |  |
| Keyword search | 44.2% |  | 79.1% | 82.6% | 84.9% |
| Compared with semantic search, hybrid search gained 32 targets in the top ten and lost 1, and keyword search gained 39 and lost 1 (*P* < 10<sup>−6</sup> for both) |  |  |  |  |  |

\*Over all 332 held-out targets with a term, hybrid search placed 70.2% first, 89.5% in the top five, 92.8% in the top ten, 94.9% in the top twenty and 96.7% in the top fifty. These values count 4 searches as misses. Hybrid search declined 3 of them, because their terms (ARS, Nodal and Electrospray) are each in more than 20,000 records. The fourth search, for the term *isochronous* (433 records), failed on a temporary capacity error.

Semantic search placed 67.2% of the 332 targets with a term in the top ten, and 70.8% of the 48 held-out targets without one. The targets with a term were therefore not easier to find. On the identifier queries, hybrid search and keyword search did not differ significantly (70.9% against 79.1%, *P* = 0.12).

**Table S4.**
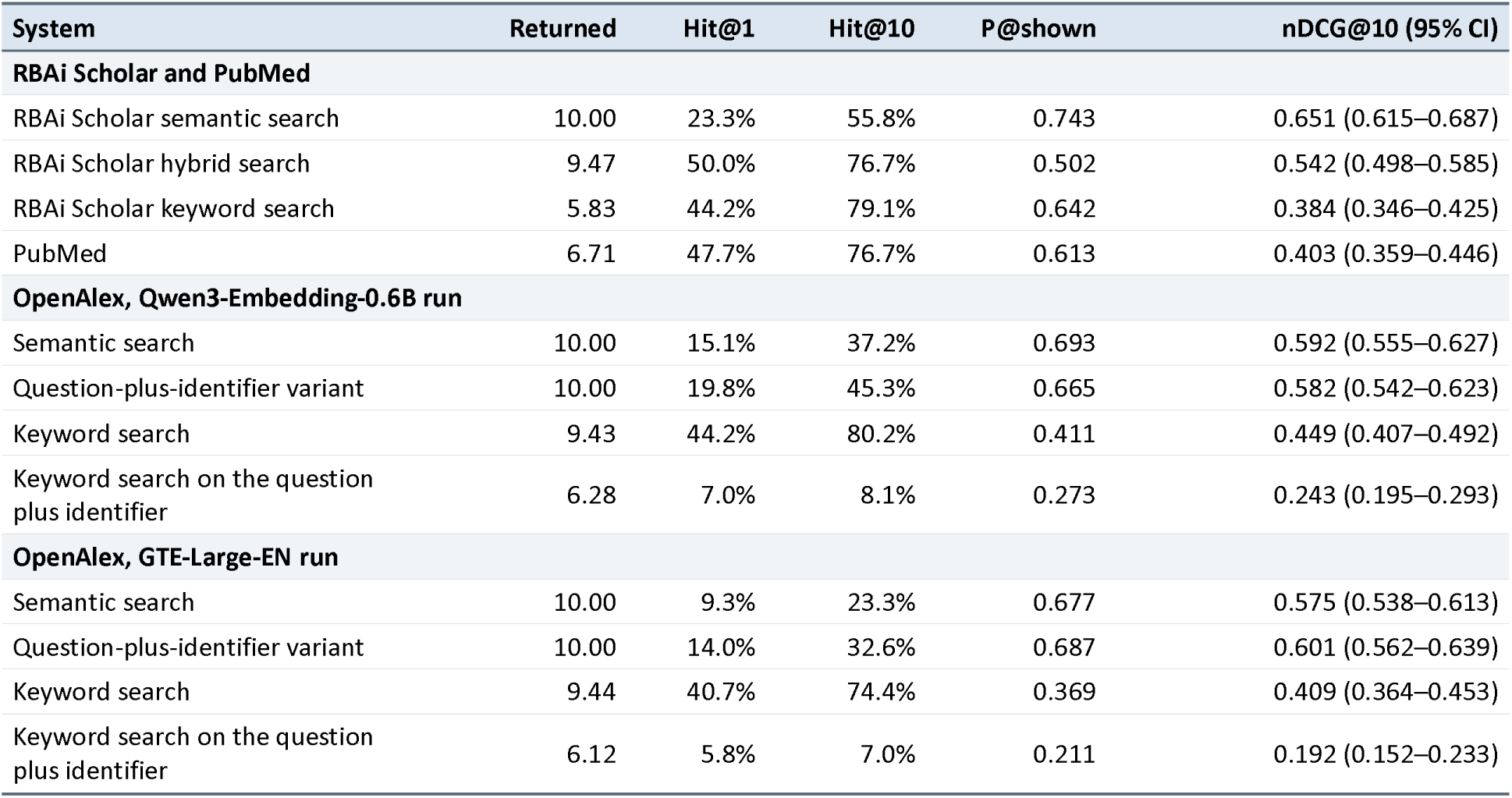
Relevance of the top ten. We asked each system for ten results on each of the 86 graded questions, and Claude Haiku 5.5 graded every result. *Returned* is the average number of results a system gave. Hit@1 and Hit@10 are the shares of questions with the target paper ranked first and in the top ten. P@shown is the share of the returned results that were graded relevant. nDCG@10 is shown with its 95% bootstrap interval. Data: Supplementary Data 1, sheet Top-ten relevance (86); results/round2-graded-2026-11-17.json.

| System | Returned | Hit@1 | Hit@10 | P@shown | nDCG@10 (95% CI) |
|---|---|---|---|---|---|
| **RBAi Scholar and PubMed** |  |  |  |  |  |
| RBAi Scholar semantic search | 10.00 | 23.3% | 55.8% | 0.743 | 0.651 (0.615–0.687) |
| RBAi Scholar hybrid search | 9.47 | 50.0% | 76.7% | 0.502 | 0.542 (0.498–0.585) |
| RBAi Scholar keyword search | 5.83 | 44.2% | 79.1% | 0.642 | 0.384 (0.346–0.425) |
| PubMed | 6.71 | 47.7% | 76.7% | 0.613 | 0.403 (0.359–0.446) |
| **OpenAlex, Qwen3-Embedding-0.6B run** |  |  |  |  |  |
| Semantic search | 10.00 | 15.1% | 37.2% | 0.693 | 0.592 (0.555–0.627) |
| Question-plus-identifier variant | 10.00 | 19.8% | 45.3% | 0.665 | 0.582 (0.542–0.623) |
| Keyword search | 9.43 | 44.2% | 80.2% | 0.411 | 0.449 (0.407–0.492) |
| Keyword search on the question plus identifier | 6.28 | 7.0% | 8.1% | 0.273 | 0.243 (0.195–0.293) |
| **OpenAlex, GTE-Large-EN run** |  |  |  |  |  |
| Semantic search | 10.00 | 9.3% | 23.3% | 0.677 | 0.575 (0.538–0.613) |
| Question-plus-identifier variant | 10.00 | 14.0% | 32.6% | 0.687 | 0.601 (0.562–0.639) |
| Keyword search | 9.44 | 40.7% | 74.4% | 0.369 | 0.409 (0.364–0.453) |
| Keyword search on the question plus identifier | 6.12 | 5.8% | 7.0% | 0.211 | 0.192 (0.152–0.233) |

**Table S5.**
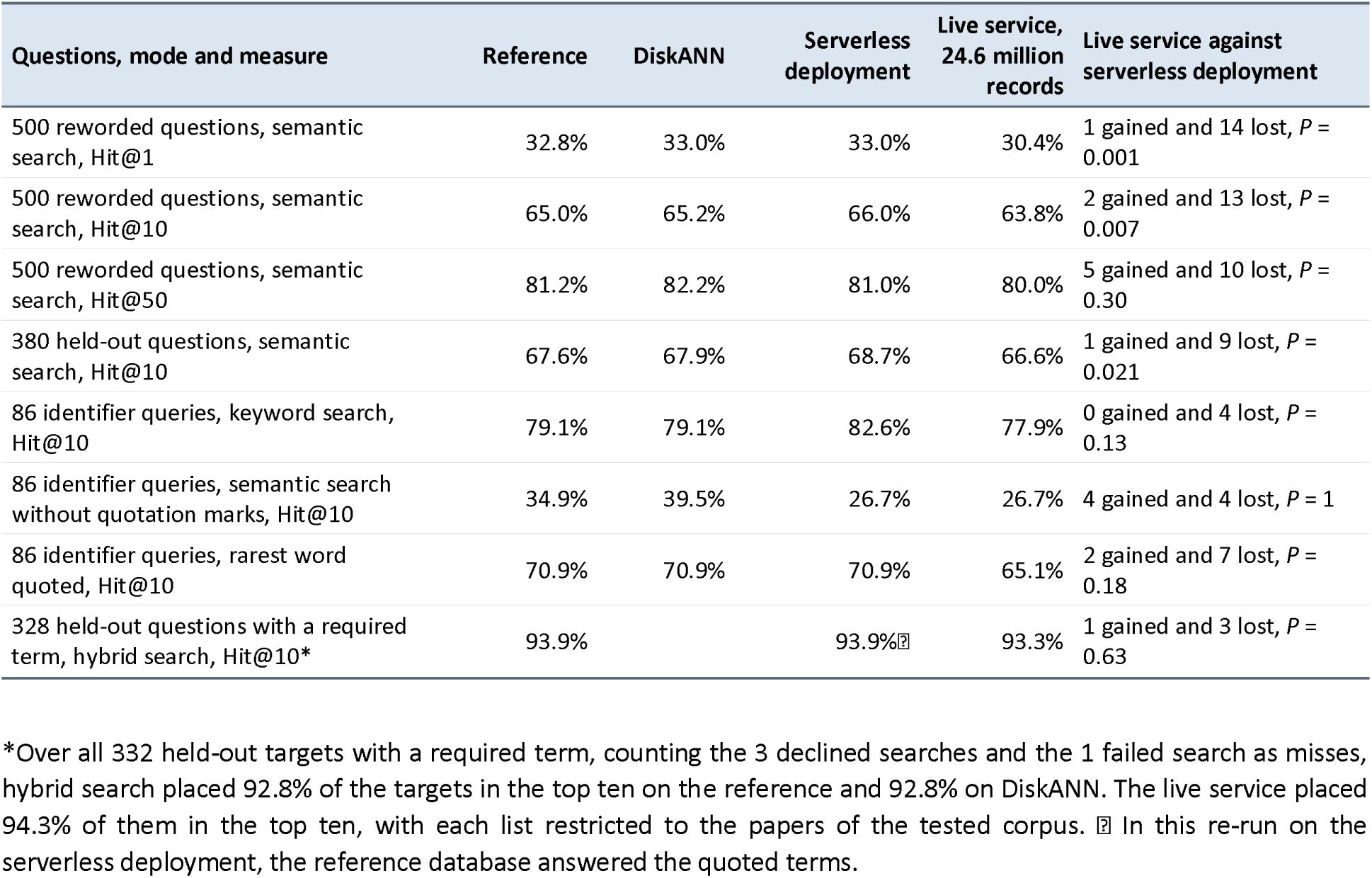
Other indexes and the live service. The table covers searches for target papers only. The reference is the implementation on which we measured every result in the main text. We measured DiskANN and the serverless deployment on the tested corpus, and the live service when it held 24.6 million records. The last column compares the live service with the serverless deployment, target by target. It counts the targets that entered the cutoff on the live service (gained) and those that left it (lost), with the exact McNemar *P*. A declined or failed search counts as a miss. Data: results/rerun-diskann.json, results/rerun-diskann-postfix.json, results/rerun-cloudflare-full.json, results/quoted-live-2026-11-14.json, results/live-24m-2026-11-16.json.

| Questions, mode and measure | Reference | DiskANN | Serverless deployment | Live service, 24.6 million records | Live service against serverless deployment |
|---|---|---|---|---|---|
| 500 reworded questions, semantic search, Hit@1 | 32.8% | 33.0% | 33.0% | 30.4% | 1 gained and 14 lost, *P* = 0.001 |
| 500 reworded questions, semantic search, Hit@10 | 65.0% | 65.2% | 66.0% | 63.8% | 2 gained and 13 lost, *P* = 0.007 |
| 500 reworded questions, semantic search, Hit@50 | 81.2% | 82.2% | 81.0% | 80.0% | 5 gained and 10 lost, *P* = 0.30 |
| 380 held-out questions, semantic search, Hit@10 | 67.6% | 67.9% | 68.7% | 66.6% | 1 gained and 9 lost, *P* = 0.021 |
| 86 identifier queries, keyword search, Hit@10 | 79.1% | 79.1% | 82.6% | 77.9% | 0 gained and 4 lost, *P* = 0.13 |
| 86 identifier queries, semantic search without quotation marks, Hit@10 | 34.9% | 39.5% | 26.7% | 26.7% | 4 gained and 4 lost, *P* = 1 |
| 86 identifier queries, rarest word quoted, Hit@10 | 70.9% | 70.9% | 70.9% | 65.1% | 2 gained and 7 lost, *P* = 0.18 |
| 328 held-out questions with a required term, hybrid search, Hit@10* | 93.9% |  | 93.9%[illegible] | 93.3% | 1 gained and 3 lost, *P* = 0.63 |

\*Over all 332 held-out targets with a required term, counting the 3 declined searches and the 1 failed search as misses, hybrid search placed 92.8% of the targets in the top ten on the reference and 92.8% on DiskANN. The live service placed 94.3% of them in the top ten, with each list restricted to the papers of the tested corpus. [illegible] In this re-run on the serverless deployment, the reference database answered the quoted terms.

The live service also kept its lead over OpenAlex semantic search with Qwen3-Embedding-0.6B. It placed 95 targets in its top ten that OpenAlex missed, and OpenAlex placed 5 that it missed (*P* < 10⁻⁶; results/openalex-500-2026-11-17.json).

**Table S6.** Re-ranking with MedCPT’s cross-encoder. MedCPT’s cross-encoder (Jin *et al*., 2023) re-ordered the same candidates that RBAi Scholar returned for each query, so the two orderings differ only in how they score those candidates. The two counts are the targets that only the cross-encoder, and only RBAi Scholar’s ranking, placed first (exact McNemar test). Data: results/medcpt-500.json, results/medcpt-round2.json.

| Queries | RBAi Scholar's ranking, Hit@1 | MedCPT cross-encoder, Hit@1 | Gained and lost at rank 1 |
|---|---|---|---|
| 500 reworded questions | 32.8% | 48.6% | 113 gained and 34 lost, *P* < 10<sup>−6</sup> |
| 86 identifier queries | 44.2% | 58.1% | 13 gained and 1 lost, *P* = 0.0018 |
| 328 questions with a required term | 71.0% | 78.0% | 53 gained and 30 lost, *P* = 0.015 |

The cross-encoder scores each query and candidate together, so it must run once for every candidate of every query.

## References

Azzopardi L. et al. (2007) Building simulated queries for known-item topics: an analysis using six European languages. In: Proceedings of the 30th Annual International ACM SIGIR Conference on Research and Development in Information Retrieval (SIGIR ‘07). ACM, pp. 455–462. doi:10.1145/1277741.1277820.

Cormack G.V. et al. (2009) Reciprocal rank fusion outperforms Condorcet and individual rank learning methods. In: Proceedings of SIGIR 2009. ACM, pp. 758–759.

Fiorini N. et al. (2018) Best Match: new relevance search for PubMed. PLoS Biol., 16, e2005343.

Järvelin K. and Kekäläinen J. (2002) Cumulated gain-based evaluation of IR techniques. ACM Trans. Inf. Syst., 20, 422–446.

Jin Q. et al. (2023) MedCPT: contrastive pre-trained transformers with large-scale PubMed search logs for zero-shot biomedical information retrieval. Bioinformatics, 39, btad651.

Malkov Y.A. and Yashunin D.A. (2020) Efficient and robust approximate nearest neighbor search using hierarchical navigable small world graphs. IEEE Trans. Pattern Anal. Mach. Intell., 42, 824–836.

Model Context Protocol (2025) Model Context Protocol specification. https://modelcontextprotocol.io/ (accessed 12 November 2026).

OpenAlex (2026) Semantic search. OpenAlex help pages. https://help.openalex.org/api/semantic-search/ (last updated 3 November 2026; accessed 24 November 2026).

Priem J. (2026) Semantic search just got a lot better, especially in non-English. openalex-users mailing list, 23 September 2026. https://groups.google.com/g/openalex-users (accessed 24 November 2026).

Priem J. et al. (2022) OpenAlex: a fully open index of scholarly works, authors, venues, institutions and concepts. arXiv:2205.01833.

Robertson S. and Zaragoza H. (2009) The probabilistic relevance framework: BM25 and beyond. Found. Trends Inf. Retr., 3, 333–389.

Sciavolino C. et al. (2021) Simple entity-centric questions challenge dense retrievers. In: *Proceedings of EMNLP* 2021. ACL, pp. 6138–6148.

Subramanya S.J. et al. (2019) DiskANN: fast accurate billion-point nearest neighbor search on a single node. In: Advances in Neural Information Processing Systems 32 (NeurIPS 2019).

Thomas P. et al. (2024) Large language models can accurately predict searcher preferences. In: Proceedings of the 47th International ACM SIGIR Conference on Research and Development in Information Retrieval (SIGIR ’24). ACM, pp. 1930–1940.

Upadhyay S. et al. (2024) UMBRELA: UMbrela is the (open-source reproduction of the) Bing RELevance Assessor. arXiv:2406.06519.

Xu L. et al. (2025) Dense retrievers can fail on simple queries: revealing the granularity dilemma of embeddings. arXiv:2506.08592.

Yeganova L. et al. (2025) LitSense 2.0: AI-powered biomedical information retrieval with sentence and passage level knowledge discovery. Nucleic Acids Res., 53, W361–W368.

Zhang Y. et al. (2025) Qwen3 Embedding: advancing text embedding and reranking through foundation models. arXiv:2506.05176.

## Supplementary references

Cormack, G.V. et al. (2009) Reciprocal rank fusion outperforms Condorcet and individual rank learning methods. In: Proceedings of SIGIR 2009. ACM, pp. 758–759.

Järvelin, K. and Kekäläinen, J. (2002) Cumulated gain-based evaluation of IR techniques. ACM Trans. Inf. Syst., 20, 422–446.

Jin, Q. et al. (2023) MedCPT: contrastive pre-trained transformers with large-scale PubMed search logs for zero-shot biomedical information retrieval. Bioinformatics, 39, btad651.

Robertson, S. and Zaragoza, H. (2009) The probabilistic relevance framework: BM25 and beyond. Found. Trends Inf. Retr., 3, 333–389.

Zhang, Y. et al. (2025) Qwen3 Embedding: advancing text embedding and reranking through foundation models. arXiv:2506.05176.

